# b3alien: A Python package to assess the introduction rate of alien species in a FAIR and reproducible way

**DOI:** 10.1101/2025.11.17.688820

**Authors:** Maarten Trekels, Quentin Groom

## Abstract

The Kunming–Montreal Global Biodiversity Framework outlines an ambitious pathway to achieve harmony with nature by 2050, with 23 targets for 2030. Among them, Target 6 seeks to reduce invasive alien species (IAS) introductions by 50% and minimize their impacts. Achieving and monitoring progress towards this target is highly challenging, as observed first records rarely reflect true introduction rates due to detection lags influenced by survey effort, detectability, and taxonomic expertise. To address this, statistical methods, such as the approach proposed by Solow and Costello, account for detection delays and provide a more reliable basis for estimating IAS establishment rates. This forms the basis of “Headline Indicator 6.1” within a wider suite of component and complementary indicators that together describe invasion dynamics and impacts.

Reliable monitoring requires transparent, reproducible tools that can integrate diverse data sources. Here, we present b3alien, a software package developed to facilitate calculation of Target 6 indicators. Built on the biodiversity data cube framework from the Biodiversity Building Blocks for Policy (B-Cubed) project, b3alien leverages the Global Biodiversity Information Facility (GBIF) infrastructure, including its Taxonomic Backbone and the Global Register of Introduced and Invasive Species (GRIIS). By integrating GBIF occurrence records with checklists and complementary datasets, the tool enables robust estimation of IAS establishment rates while supporting additional invasion-related indicators. Outputs are technically rigorous yet accessible, ensuring usability for policymakers and stakeholders.

The basic workflow is using a GBIF-based occurrence cube, which can be extended by incorporating citizen science contributions, private datasets, and customized checklists, thereby ensuring flexibility and adaptability across contexts. By aligning with FAIR data principles, b3alien ensures indicators are findable, accessible, interoperable, and reusable. Importantly, the approach empowers countries to build their own indicators on open, community-driven infrastructures, lowering technical and financial barriers, fostering bottom-up ownership, and ensuring scientific and policy credibility.

In summary, b3alien demonstrates how open infrastructures, standardized data, and reproducible workflows can make IAS monitoring both accessible and scientifically robust. By bridging biodiversity data with actionable policy insights, it provides a practical and equitable pathway to support the ambitions of the Global Biodiversity Framework’s Target 6.

## Introduction

The Kunming-Montreal Global Biodiversity Framework (GBF), which supports the Sustainable Development Goals and builds on the Convention on Biological Diversity’s (CBD) previous Strategic Plans, sets out an ambitious pathway to reach the global vision of a world living in harmony with nature by 2050. Among the Framework’s key elements are 23 targets for 2030. Here we focus on Target 6 for invasive species “*Reduce the introduction of invasive alien species by 50% and minimize their impact*”. Despite the simple wording, this is a hugely ambitious target, not only to conduct the biosecurity and species management required to achieve it, but also to know whether progress is being made towards the target, or even whether it has been achieved. Toward that aim several indicators have been proposed, but the so-called “Headline indicator 6.1” is the rate of invasive alien species (IAS) establishment.

A major difficulty in estimating the rate of IAS establishment is that simple counts of observed first records rarely reflect true arrival rates. In most cases, there is a significant lag between when an alien species first arrives in a country, and when it is detected and reported. These lags vary depending on factors such as survey effort, intrinsic detectability of the species, taxonomic expertise, and investment in monitoring, and they can differ greatly across regions, habitats and time periods. As a result, observed records alone can lead to misleading conclusions about introduction rates. To address this issue, Solow & Costello (2004) proposed a statistical method that estimates the rate of alien species accumulation while explicitly accounting for detection lags. This approach provides a more reliable basis for quantifying introduction rates and has since been recognized as an important tool for developing robust invasion indicators (Buba et al., 2024; McGeoch et al., 2023).

To achieve any of the GBF targets, decision makers at local, regional, national, and international levels must be able to calculate indicators accurately and reliably using data sources that are both cost-effective and straightforward to collect. For Target 6, this requires comprehensive data on the status, trends, and impacts of non-native species. The resulting indicators should be presented in an actionable and accessible format, accompanied by clear measures of uncertainty and transparent provenance (Sica et al., 2024).

In this paper we present the b3alien software which was developed to address these challenges by providing a transparent, reproducible, and user-friendly approach to calculating the rate of establishment (Trekels, 2025b). Built on the concept of occurrence cubes developed in the Biodiversity Building Blocks for Policy (B-Cubed) project (Groom et al., 2023, 2025), b3alien harnesses the extensive open infrastructure of the Global Biodiversity Information Facility (GBIF). This includes access to tools such as the GBIF Taxonomic Backbone and datasets, such as the Global Register of Introduced and Invasive Species (GRIIS) (Pagad et al., 2018, 2022). By combining these resources with open biodiversity occurrence data, b3alien enables the calculation of the headline indicator for Target 6, the rate of invasive alien species establishment. Crucially, it generates outputs in formats that are both technically robust and accessible to policymakers, thus bridging the gap between raw biodiversity data and actionable policy insights.

## Description

b3alien offers a set of classes and functions to create a streamlined workflow to calculate the Target 6.1 headline indicator of the Global Biodiversity Framework. The package integrates biodiversity occurrence data with authoritative species status information, providing users with reproducible estimates of establishment rates alongside clear measures of uncertainty. Its design draws on the biodiversity data cube concept (Groom et al., 2023) and follows the indicator strategy of McGeoch et al, (2023) and UNEP-WCMC (2025). Designed for flexibility, b3alien allows workflows to be tailored to the specific reporting needs of individual countries, while ensuring consistency and reproducibility across contexts. The package is openly available on GitHub (https://github.com/mtrekels/b3alien), is fully documented through an auto-generated website (https://b3alien.readthedocs.io/), and is published on the Python Package Index (PyPI), making it straightforward to install via pip together with all dependencies. This commitment to open infrastructure and accessibility aligns with the FAIR principles, ensuring that the data and methods remain findable, accessible, interoperable, and reusable.

The workflow is designed to be straightforward and intuitive (see Fig. 1). The first step is to generate an occurrence cube of biodiversity data (Groom et al., 2023), an implementation of the concept of Essential Biodiversity Variables (Jetz et al., 2019; Pereira et al., 2013). By default, the cube is generated through the SQL API of GBIF (GBIF Secretariat, 2025), though additional filters can be applied to select particular subsets of occurrence data or include or exclude specific datasets).

**Figure 1.**
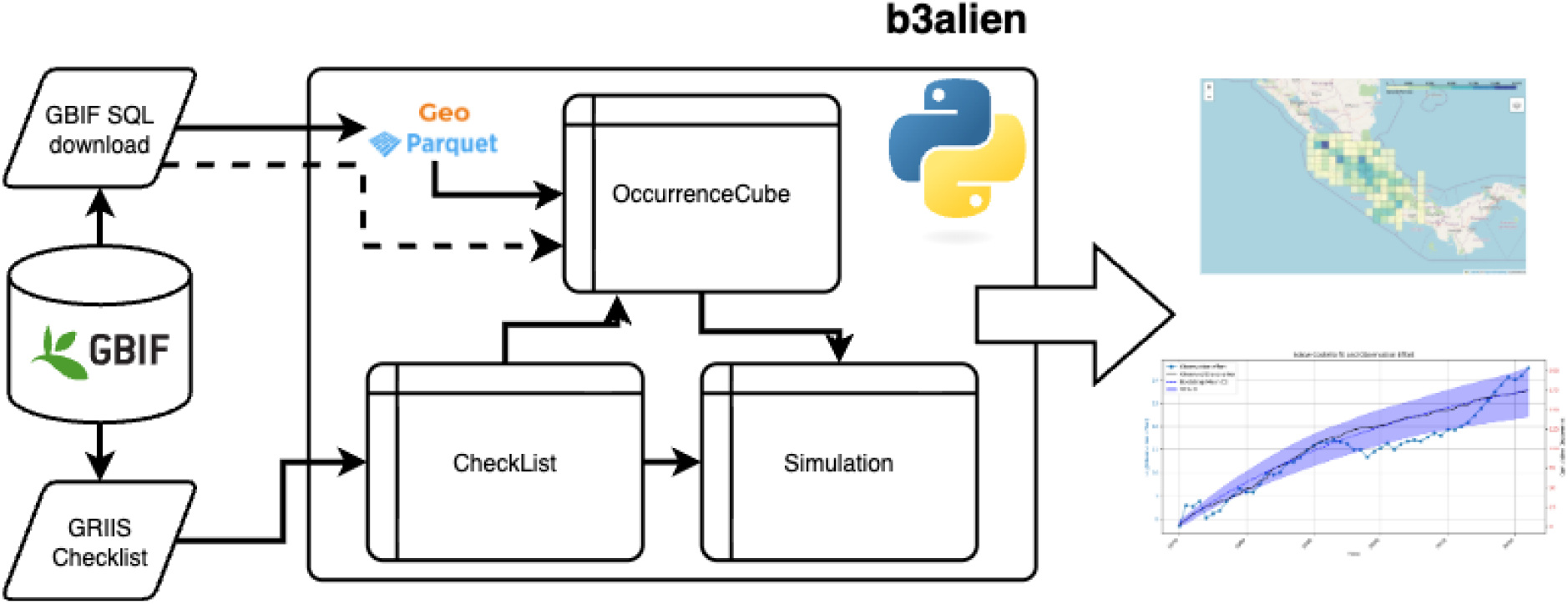
Workflow of the b3alien Python package for calculating the rate of establishment of invasive alien species (Headline indicator 6.1). Biodiversity occurrence data are retrieved from GBIF through the SQL API and stored as GeoParquet files to build an occurrence cube. If no spatial information is needed, the generation of the GeoParquet file can be bypassed. Species status information is obtained from national GRIIS checklists and harmonized with the GBIF Taxonomic Backbone. These inputs feed into the Simulation module, which applies the Solow–Costello algorithm to estimate establishment rates with confidence intervals.

A step by step guide to creating the occurrence cube is included in Appendix A. The Target 6.1 indicator can be calculated by using a standard GBIF cube, but the SQL statement can be expanded to include a measure of the observation effort. The standard workflow also allows inclusion of proxies for observation effort, such as the number of distinct observers (dwc:recordedBy). These can be assessed spatially by grid cell (see Fig. 2b) or through time (see Fig. 3), thereby providing insights into variation in survey intensity., However, this part of the workflow can be replaced by more sophisticated approaches in case other data are available on the geographic area under consideration (see Leihy et al., (2025)). Outputs include spatial and temporal visualizations that can be linked to other environmental datasets.

**Figure 2.**
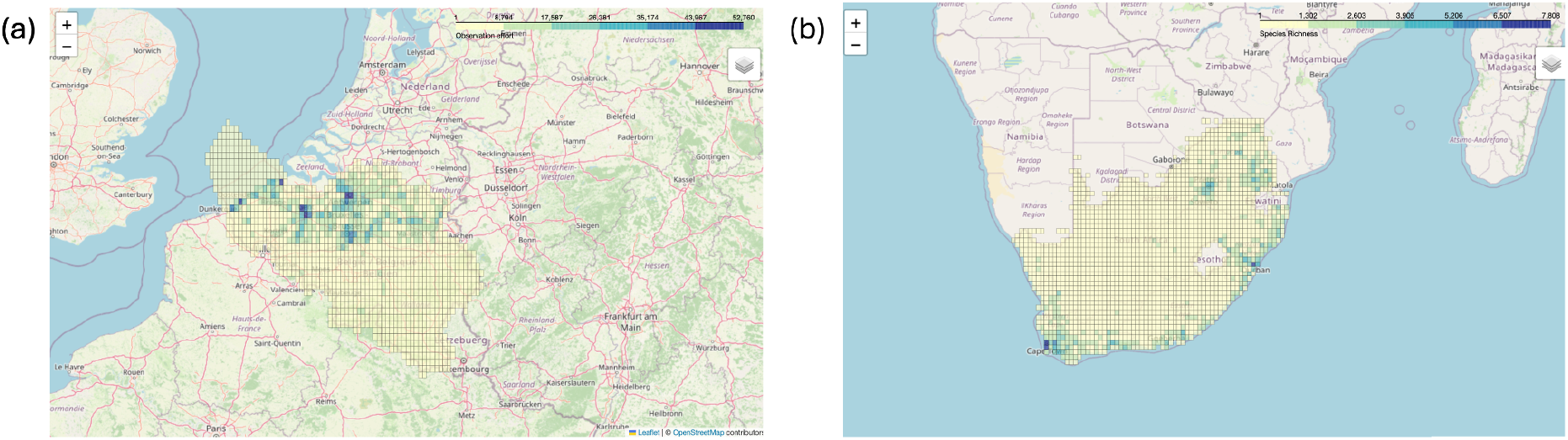
Interactive visualization of biodiversity data cubes using the b3alien package. (a) Observed species richness in Belgium. (b) Number of distinct observers per grid cell in South Africa. Both maps aggregate GBIF occurrence records at level-2 Quarter Degree Grid Cells, providing geospatial context that supports interpretation of species distributions and observation effort.

**Figure 3.**
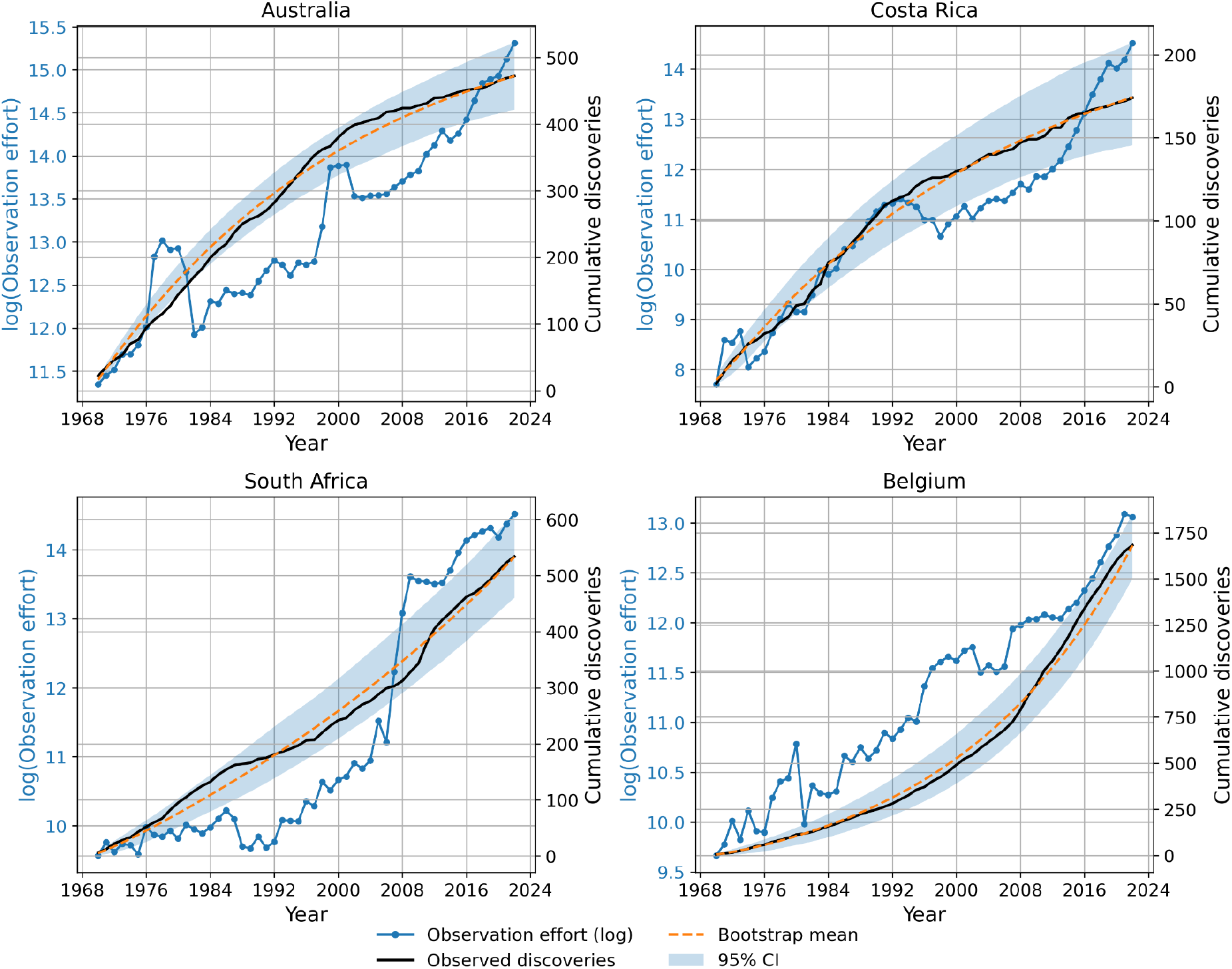
Observed and simulated cumulative number of introduced species for Belgium, South Africa, Australia and Costa Rica. Black lines show the observed number of introductions from the GBIF data cube; Dashed lines are simulations fitted with the Solow-Costello model; Dotted line represents the logarithm of the number of distinct observers in the data cube.

To decide on whether a species is native or introduced, information per country on the species is determined through the GRIIS checklists (Pagad et al., 2018, 2022). Most national lists are published as simple species–status pairs, in which case b3alien provides functions to map names onto the GBIF Taxonomic Backbone for harmonization. In this case, the b3alien package provides functions to map the species names to the GBIF taxonomic backbone. This matching is needed to be able to match the species in the data cube with the GRIIS checklist. In case the information in the checklist is only present at genus level, all species within that genus will be taken into account in the calculation of the cumulative number of species.

This method assumes that species not contained in the GRIIS checklist are native species. This however is in most cases a simplification, and potentially doesn’t cover all alien species. Using b3alien, species can be added to or removed from the CheckList object, by using the dwc:taxonID field of the GBIF taxonomic backbone. This allows the users to review their GRIIS list and use the most up-to-date available information.

In rare cases, such as Belgium (Reyserhove et al., 2020), checklists contain expert-assessed introduction dates, which allow direct estimation of establishment rates without recourse to GBIF occurrence data. This is the ideal situation, where it is not necessary to combine the checklist with GBIF occurrence data. The software package can calculate the rate of instruction directly from the GRIIS checklist.

In the following sections, we describe how the package leverages biodiversity occurence cubes. Second, we present the main application of the package, estimating the rate of establishment of invasive alien species using the Solow–Costello method.

## Biodiversity occurrence cubes

Although the primary purpose of the b3alien Python package is to calculate the rate of establishment, the use of biodiversity occurrence cubes as an input format provides several additional features. In its simplest form, a GBIF cube contains only the text identifier of each grid cell, and the workflow runs without issue on this basic structure.

If a GeoPackage (Open Geospatial Consortium, 2024b) file exists for the used grid system (e.g. Larsen 2021), the utility functions inside the b3alien package can create a GeoParquet file (Open Geospatial Consortium, 2024a). This file format is particularly well suited for tabular data linked to vector geometries. It stores geometry in Well-Known Binary (WKB) format, which reduces overall file size compared to the original data. An example of a GeoParquet file is provided in Trekels (2025a).

Using a geospatial format makes it possible to plot the data on maps and overlay it with remote-sensing products. When the b3alien package is run in a Jupyter notebook environment, it automatically employs Folium (Folium Developers, 2025) to visualize the data. Fig. 2 illustrates this with two examples (a) “observed species richness” in Belgium (all taxa) and (b) number of distinct observers in South Africa. Clear, easily interpretable visualizations of the data can support informed decision making, especially when biodiversity data are spatially heterogeneous.

By storing the cube in standard geospatial formats and representing it in both Xarray and DataFrame structures, the calculation of additional indicators becomes straightforward. These widely used Python formats support simple yet flexible operations, such as filtering, aggregation and rescaling, that can be applied directly to the cube to generate new outputs.

The software also supports loading GeoParquet files directly from Google Cloud services, enabling the easy reuse of biodiversity data cubes for future analyses. These cubes can be integrated into other platforms, such as Google Earth Engine, to broaden their applications. Support for other cloud providers can be envisaged if there is sufficient demand.

## Rate of Establishment

The main purpose of this package is to calculate the rate of establishment and the change of it. In order to showcase the calculation of this indicator, we developed a Jupyter notebook (Appendix B) that was run for four different data cubes generated for Belgium (GBIF.Org User, 2025), Australia (GBIF.org User, 2025a), Costa Rica (GBIF.org User, 2025b) and South Africa (GBIF.org User, 2025c). The cumulative number of alien species is shown in Fig. 3, including the fit according to the Solow-Costello model. The determination of the confidence intervals on the rate of establishment is computationally intensive and this calculation is parallelized using all but one core of the machine it is running on. The final results of the calculations are shown in Table 1.

**Table 1:**
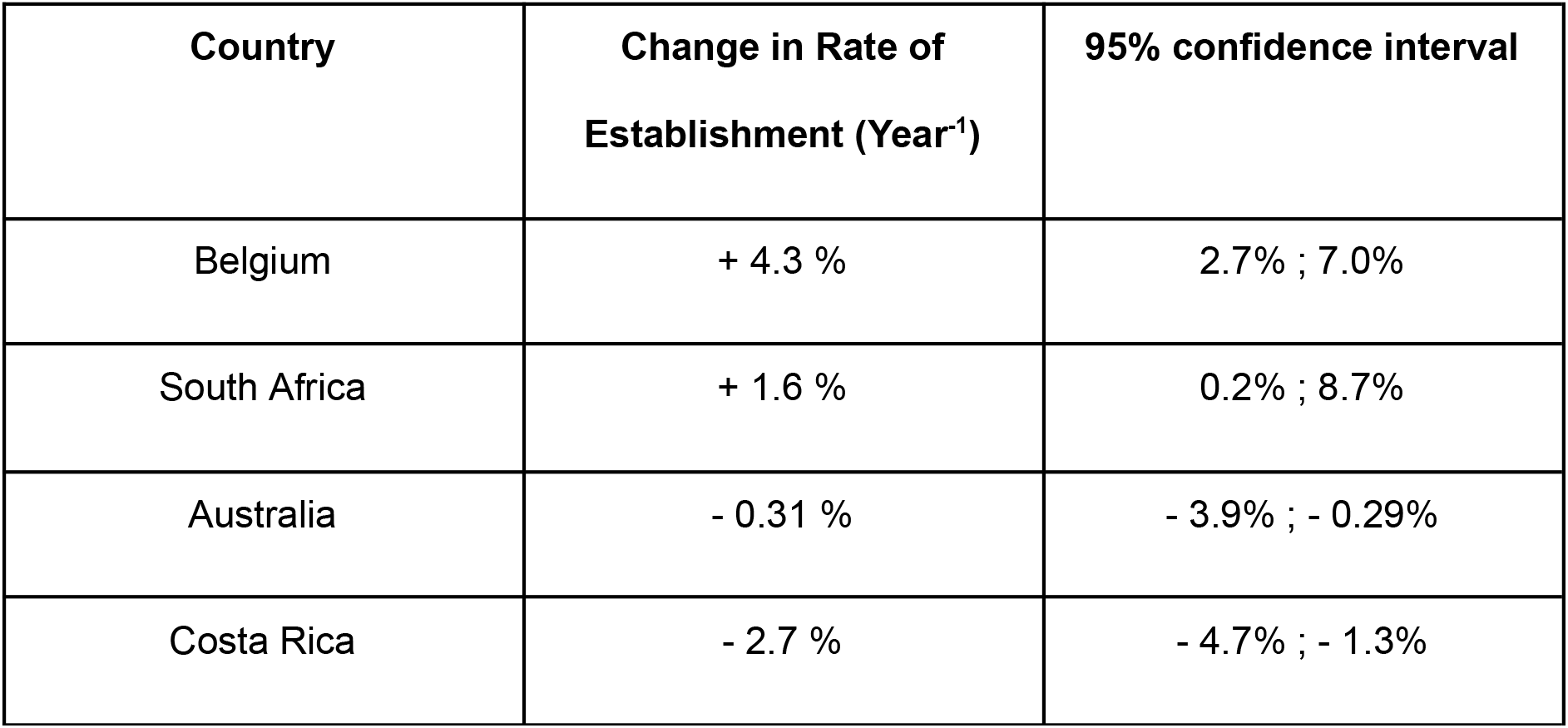
Estimated annual change in the rates of establishment of invasive alien species calculated from GBIF occurrence data for four countries. Values represent percentage change per year, with 95% confidence intervals shown in brackets.

In order to cross validate the results on the rate of introduction, the detailed GRIIS checklist of Belgium was used (Desmet et al., 2025). Since this checklist contains an expert evaluation of the date of introduction of species, it was possible to run the Solow-Costello simulation on this detailed dataset.

The change in introduction rate calculated from the detailed GRIIS checklist is +6.5% CI:[3.8%; 7.9%] for Belgium, compared to +4.5% CI:[2.7%; 7.0%] calculated for the GBIF datacube. This shows that there might be an underestimation of the rate based on GBIF occurrences. This can be due to multiple factors, such as missing observations that were accounted for in the checklist. A lag in publishing observation data is well known (Christie et al., 2021). Although the point estimates differ, their 95% confidence intervals overlap, suggesting no strong evidence for a significant difference..

## Discussion

Although repeatable and user-friendly methods for calculating these indicators are essential, even the wealthiest countries often struggle to produce them reliably (Buchanan et al., 2020). This highlights the need to simplify the process and ensure that indicators are accessible to all. Enabling countries to take ownership of their own biodiversity assessments in a bottom-up manner fosters responsible local management, rather than creating the perception of a top-down framework imposed by richer nations (Navarro et al., 2017; Secretariat of the Convention on Biological Diversity, 2020).

Starting the workflow with generating a GBIF occurrence cube, allows for the flexibility of including or excluding specific occurrences or datasets in the calculation. This can be used to examine the impact of particular aspects of the data, such as the contribution of citizen science platforms (such as iNaturalist or Observations.org), or to define other measures for observation effort and bias. Moreover, additional aggregated parameters from the GBIF data can be included as an additional dimension in the data cube. Note that it is not mandatory to use only the data published on GBIF. Once a GBIF cube is generated, other (private) biodiversity data sets can be included in the data cube as long as the data are georeferenced. This would only require a minor modification of the current workflow. As stated above, it is also possible to add additional species to the GRIIS checklist in case of data gaps.

With b3alien, we developed a software package that provides an easy and reproducible workflow for reporting on the Target 6 indicators of the Global Biodiversity Framework. The workflow is transferable to any country and can be used to estimate the rate of alien species introductions. In particular, when no detailed checklist is available, it offers an efficient way to infer introduction dates from GBIF occurrence data. By applying this workflow, the calculation of Target 6 indicators becomes transparent, consistent, and straightforward to replicate. Importantly, it also embodies the FAIR principles, ensuring that biodiversity data and derived indicators are findable, accessible, interoperable, and reusable, which was precisely the aim of promoting FAIR data practices in biodiversity monitoring and policy (Sica et al., 2024).

This bottom-up approach is reinforced by relying on open, community-driven infrastructures, which give countries the tools to build their own indicators on a foundation that is both globally consistent and locally adaptable. An additional strength of b3alien is that it builds on the well-established open infrastructure of GBIF. This ensures long-term sustainability, taxonomic consistency through the GBIF Taxonomic Backbone, and authoritative species lists via resources such as GRIIS. As these infrastructures are community-driven and continuously updated, indicators derived from b3alien benefit from global data sharing while avoiding duplication of effort. This reliance on transparent and trusted infrastructures increases both the scientific robustness and the policy credibility of the results..

Moreover, the use of occurrence cubes allows for easy visualisation of these results in relation to other geospatial products. This can be for educational purposes, but also allows to relate the results from the calculations to other environmental variables. This facilitates the calculation of the component and complementary indicators, since the spatial component as well as the normalization tools allow the user to calculate additional parameters of a biological invasion, such as the rate of spread of a species.

In summary, b3alien demonstrates how open infrastructures, standardized datasets, and reproducible workflows can make the monitoring of invasive alien species both accessible and scientifically robust. By aligning with the FAIR principles and leveraging community-driven resources such as GBIF and GRIIS, it enables transparent and comparable calculation of the Target 6 indicators across countries. This not only lowers technical and financial barriers but also empowers countries to take ownership of their biodiversity reporting in a bottom-up manner. Ultimately, the approach offers a practical pathway towards meeting the ambitions of the Kunming–Montreal Global Biodiversity Framework by providing decision makers with credible, actionable, and equitable measures of progress.

## Author contributions

MT and QG conceived the study. MT curated the data, carried out the formal analyses, developed the methodology, implemented the software, validated results, and produced the visualizations. QG acquired funding, provided project administration, and supervised the work. Both authors contributed to resources. MT and QG wrote the original draft and both authors reviewed and edited the manuscript.

## Acknowledgements

This software was developed under the B3 project. B3 (Biodiversity Building Blocks for policy) receives funding from the European Union’s Horizon Europe Research and Innovation Programme (ID No 101059592). Views and opinions expressed are however those of the author(s) only and do not necessarily reflect those of the European Union or the European Commission. Neither the EU nor the EC can be held responsible for them.

## Notes

https://doi.org/10.15468/DL.WCNHE7

https://doi.org/10.15468/DL.M2B4CR

https://doi.org/10.15468/DL.CDZKWX

https://doi.org/10.15468/DL.PHSWQQ

https://doi.org/10.5281/ZENODO.15738139

### Competing Interest Statement

The authors have declared no competing interest.

## References

Buba, Y., Kiflawi, M., McGeoch, M. A., & Belmaker, J. (2024). Evaluating models for estimating introduction rates of alien species from discovery records. Global Ecology and Biogeography, 33(8), e13859. 10.1111/geb.13859

Buchanan, G. M., Butchart, S. H. M., Chandler, G., & Gregory, R. D. (2020). Assessment of national-level progress towards elements of the Aichi Biodiversity Targets. Ecological Indicators, 116, 106497. 10.1016/j.ecolind.2020.106497

Christie, A. P., White, T. B., Martin, P. A., Petrovan, S. O., Bladon, A. J., Bowkett, A. E., Littlewood, N. A., Mupepele, A.-C., Rocha, R., Sainsbury, K. A., Smith, R. K., Taylor, N. G., & Sutherland, W. J. (2021). Reducing publication delay to improve the efficiency and impact of conservation science. PeerJ, 9, e12245. 10.7717/peerj.12245

Desmet, P., Reyserhove, L., Oldoni, D., Groom, Q., Adriaens, T., Vanderhoeven, S., & Shyama Pagad. (2025). Global Register of Introduced and Invasive Species—Belgium [Dataset]. Invasive Species Specialist Group ISSG. 10.15468/XOIDMD

Folium Developers. (2025). Folium (Version 0.20.0) [Computer software]. https://python-visualization.github.io/folium/latest/

GBIF Secretariat. (2025). API SQL Downloads. API SQL Downloads:: Technical Documentation. https://techdocs.gbif.org/en/data-use/api-sql-downloads

GBIF.org User. (2025a). Occurrence Download Australia (p. 1033250931) [Text/tab-separated-values,application/zip]. The Global Biodiversity Information Facility. 10.15468/DL.WCNHE7

GBIF.Org User. (2025). Occurrence Download Belgium (p. 194091876) [Text/tab-separated-values,application/zip]. The Global Biodiversity Information Facility. 10.15468/DL.M2B4CR

GBIF.org User. (2025b). Occurrence Download Costa Rica (p. 111100518) [Text/tab-separated-values,application/zip]. The Global Biodiversity Information Facility. 10.15468/DL.CDZKWX

GBIF.org User. (2025c). Occurrence Download South Africa (p. 458796333) [Text/tab-separated-values,application/zip]. The Global Biodiversity Information Facility. 10.15468/DL.PHSWQQ

Groom, Q., Abraham, L., Adriaens, T., Breugelmans, L., Clarke, D. A., Di Musciano, M., Dove, S., Estupinan‐Suarez, L. M., Faulkner, K. T., Fernandez, M., Hendrickx, L. A., Hui, C., Joly, A., Kumschick, S., Langeraert, W., Martini, M., Miller, J., Oldoni, D., Pereira, H., … Desmet, P. (2025). Creating the vision of rapid, repeatable, reactive data workflows for policy on biodiversity. Ecological Solutions and Evidence, 6(3), e70113. 10.1002/2688-8319.70113

Groom, Q., Abraham, L., Adriaens, T., Breugelmans, L., Clarke, D., Fernández, M., Hendrickx, L., Hui, C., Kumschick, S., Martini, M., McGeoch, M., Metodiev, T., Miller, J., Oldoni, D., Pereira, H., Preda, C., Robertson, T., Rocchini, D., Seebens, H., … Desmet, P. (2023). B-Cubed: Leveraging Analysis-Ready Biodiversity Datasets and Cloud Computing for Timely and Actionable Biodiversity Monitoring. Biodiversity Information Science and Standards, 7, e110734. 10.3897/biss.7.110734

Jetz, W., McGeoch, M. A., Guralnick, R., Ferrier, S., Beck, J., Costello, M. J., Fernandez, M., Geller, G. N., Keil, P., Merow, C., Meyer, C., Muller-Karger, F. E., Pereira, H. M., Regan, E. C., Schmeller, D. S., & Turak, E. (2019). Essential biodiversity variables for mapping and monitoring species populations. Nature Ecology & Evolution, 3(4), 539–551. 10.1038/s41559-019-0826-1

Larsen, R. (2021). Qdgc South Africa (Version 1.0.0) [Dataset]. Zenodo. 10.5281/ZENODO.4457815

Leihy, R. I., McGeoch, M. A., Clarke, D. A., Peake, L., Buba, Y., Belmaker, J., & Chown, S. L. (2025). Antarctic Biosecurity Policy Effectively Manages the Rates of Alien Introductions. Earth’s Future, 13(4), e2024EF005405. 10.1029/2024EF005405

McGeoch, M. A., Buba, Y., Arlé, E., Belmaker, J., Clarke, D. A., Jetz, W., Li, R., Seebens, H., Essl, F., Groom, Q., García‐Berthou, E., Lenzner, B., Meyer, C., Vicente, J. R., Wilson, J. R. U., & Winter, M. (2023). Invasion trends: An interpretable measure of change is needed to support policy targets. Conservation Letters, 16(6), e12981. 10.1111/conl.12981

Navarro, L. M., Fernández, N., Guerra, C., Guralnick, R., Kissling, W. D., Londoño, M. C., Muller-Karger, F., Turak, E., Balvanera, P., Costello, M. J., Delavaud, A., El Serafy, G., Ferrier, S., Geijzendorffer, I., Geller, G. N., Jetz, W., Kim, E.-S., Kim, H., Martin, C. S., … Pereira, H. M. (2017). Monitoring biodiversity change through effective global coordination. Current Opinion in Environmental Sustainability, 29, 158–169. 10.1016/j.cosust.2018.02.005

Open Geospatial Consortium. (2024a). Geoparquet (Version v1.1.0) [Computer software]. https://github.com/opengeospatial/geoparquet

Open Geospatial Consortium. (2024b). GeoPackage (Version 1.4.0). http://www.opengis.net/doc/IS/geopackage/1.4

Pagad, S., Bisset, S., Genovesi, P., Groom, Q., Hirsch, T., Jetz, W., Ranipeta, A., Schigel, D., Sica, Y. V., & McGeoch, M. A. (2022). Country Compendium of the Global Register of Introduced and Invasive Species. Scientific Data, 9(1), 391. 10.1038/s41597-022-01514-z

Pagad, S., Genovesi, P., Carnevali, L., Schigel, D., & McGeoch, M. A. (2018). Introducing the Global Register of Introduced and Invasive Species. Scientific Data, 5(1), 170202. 10.1038/sdata.2017.202

Pereira, H. M., Ferrier, S., Walters, M., Geller, G. N., Jongman, R. H. G., Scholes, R. J., Bruford, M. W., Brummitt, N., Butchart, S. H. M., Cardoso, A. C., Coops, N. C., Dulloo, E., Faith, D. P., Freyhof, J., Gregory, R. D., Heip, C., Höft, R., Hurtt, G., Jetz, W., … Wegmann, M. (2013). Essential Biodiversity Variables. Science, 339(6117), 277–278. 10.1126/science.1229931

Reyserhove, L., Desmet, P., Oldoni, D., Adriaens, T., Strubbe, D., Davis, A. J. S., Vanderhoeven, S., Verloove, F., & Groom, Q. (2020). A checklist recipe: Making species data open and FAIR. Database, 2020, baaa084. 10.1093/database/baaa084

Secretariat of the Convention on Biological Diversity. (2020). Global Biodiversity Outlook 5. https://www.cbd.int/gbo/gbo5/publication/gbo-5-en.pdf

Sica, Y., Seebens, H., Fernandez, M., Hughes, A. C., Cang, H., Kumschick, S., Estupinan-Suarez, L. M., Groom, Q., Niamir, A., Gudde, R., Krug, R. M., Hendrickx, L., Yovcheva, N., Gill, M. J., Rodrigues, A., & Gonzalez, A. (2024). Effective biodiversity monitoring requires FAIR data and FAIR models for FAIR indicators (Findable, Accessible, Interoperable, and Reusable) [Policy Brief]. 10.5281/ZENODO.13912947

Solow, A. R., & Costello, C. J. (2004). Estimating the rate of species introductions from the discovery records. Ecology, 85(7), 1822–1825. 10.1890/03-3102

Trekels, M. (2025a). Biodiversity Data Cube of South Africa at level 2 EQDGC [Dataset]. Zenodo. 10.5281/ZENODO.15738139

Trekels, M. (2025b). b3alien: A Python Package to Calculate the Global Biodiversity Framework Target 6 indicators (Version 0.2.2) [Computer software]. Zenodo. 10.5281/ZENODO.17054388

UNEP-WCMC. (2025). Indicators for the Kunming – Montreal Global Biodiversity Framework. https://gbf-indicators.org/

