## Appendix A for "b3alien: A Python package to assess the introduction rate of alien species in a FAIR and reproducible way"

APPENDIX A: Creating a GBIF Occurrence Cube

In this appendix we explain how you create a GBIF occurrence cube. Both by using the graphical user interface and the SQL API of GBIF.

### Using the graphical user interface

The easiest way to generate a data cube is through the graphical user interface. The detailed instructions on how to create a data cube are explained in the following tutorial [1]: <https://docs.b-cubed.eu/tutorials/download-a-cube-from-gbif/>

Currently the workflow is designed to work with monthly data, which means that you should select ‘Year-Month’ as the time dimension.

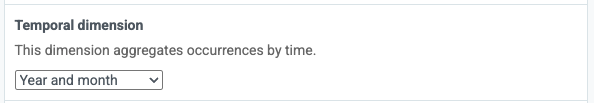

Before the download of the data cube starts, the user will see an overview of the SQL command that is used to generate the cube (see figure 1).

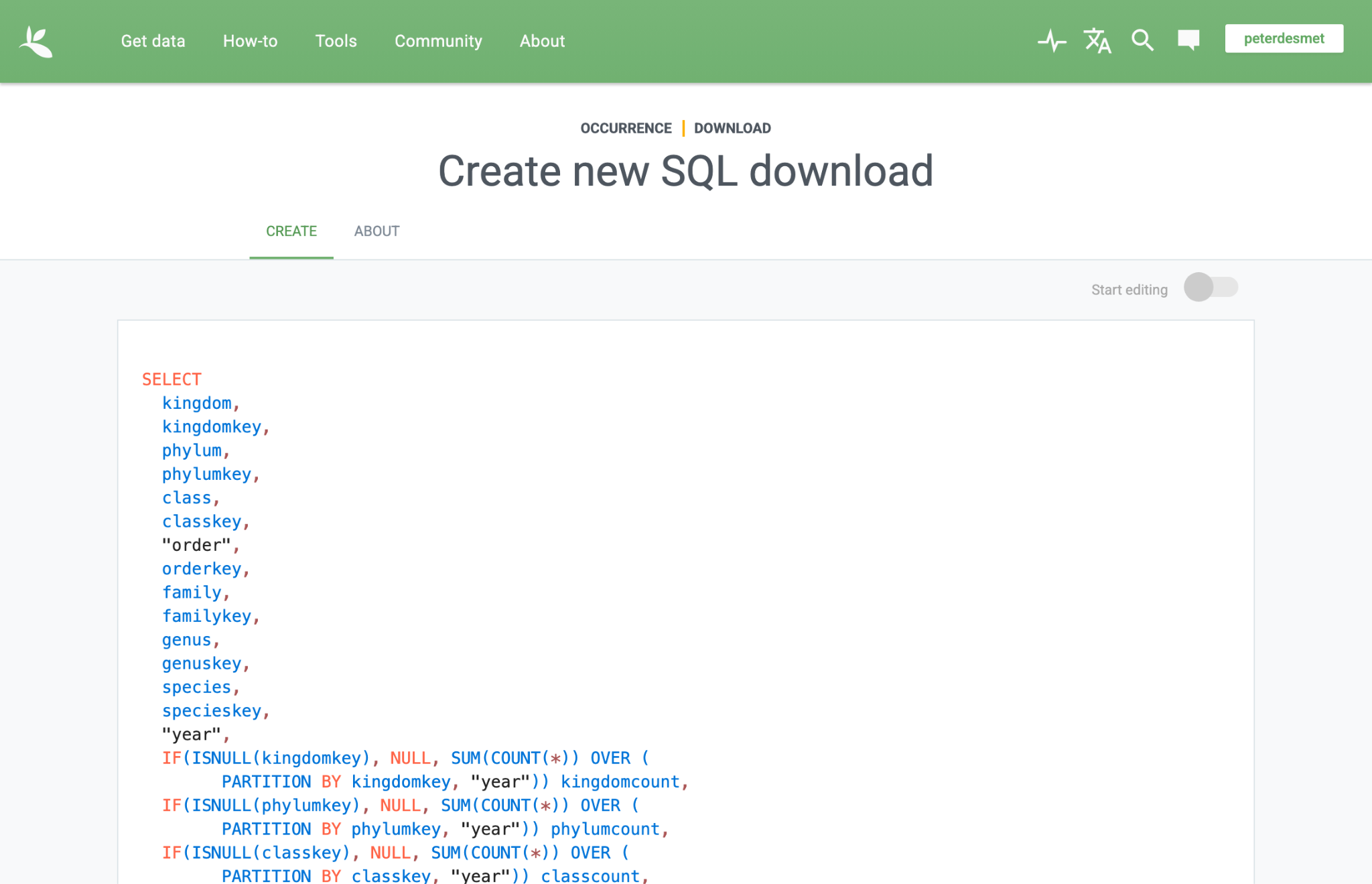

Figure 1: SQL interface that the user will see when generating the biodiversity data cube.

In case additional dimensions are wanted (e.g. to select the number of distinct observers per grid cell), the user can modify the SQL command by clicking the “Start editing” slider on the top right corner of the SQL statement.

### Using the GBIF SQL API

However, it is not necessary to go through the graphical user interface. The SQL statement can also directly be sent to the GBIF api. A detailed description of the GBIF API can be found in [2].

The statement that is used in the exemplar workflow of Appendix B, is the following:

SELECT

kingdom,

kingdomkey,

phylum,

phylumkey,

class,

classkey,

"order",

orderkey,

family,

familykey,

genus,

genuskey,

species,

specieskey,

PRINTF('%04d-%02d', "year", "month") yearmonth,

GBIF_EQDGCODE(2, decimallatitude, decimallongitude, 0.0) eqdcellcode,

IF(ISNULL(kingdomkey), NULL, SUM(COUNT(*)) OVER (

PARTITION BY kingdomkey, GBIF_EQDGCODE(2, decimallatitude, decimallongitude, 0.0), PRINTF('%04d-%02d', "year", "month"))) kingdomcount,

IF(ISNULL(phylumkey), NULL, SUM(COUNT(*)) OVER (

PARTITION BY phylumkey, GBIF_EQDGCODE(2, decimallatitude, decimallongitude, 0.0), PRINTF('%04d-%02d', "year", "month"))) phylumcount,

IF(ISNULL(classkey), NULL, SUM(COUNT(*)) OVER (

PARTITION BY classkey, GBIF_EQDGCODE(2, decimallatitude, decimallongitude, 0.0), PRINTF('%04d-%02d', "year", "month"))) classcount,

IF(ISNULL(familykey), NULL, SUM(COUNT(*)) OVER (

PARTITION BY familykey, GBIF_EQDGCODE(2, decimallatitude, decimallongitude, 0.0), PRINTF('%04d-%02d', "year", "month"))) familycount,

IF(ISNULL(genuskey), NULL, SUM(COUNT(*)) OVER (

PARTITION BY genuskey, GBIF_EQDGCODE(2, decimallatitude, decimallongitude, 0.0), PRINTF('%04d-%02d', "year", "month"))) genuscount,

IF(ISNULL(orderkey), NULL, SUM(COUNT(*)) OVER (

PARTITION BY orderkey, GBIF_EQDGCODE(2, decimallatitude, decimallongitude, 0.0), PRINTF('%04d-%02d', "year", "month"))) ordercount,

COUNT(DISTINCT recordedby) distinctobservers,

COUNT(*) occurrences

FROM

occurrence

WHERE

occurrence.countrycode = 'CR'

AND (occurrence.level0gid = 'CRI' OR occurrence.level1gid = 'CRI' OR occurrence.level2gid = 'CRI' OR occurrence.level3gid = 'CRI')

AND occurrence.occurrencestatus = 'PRESENT'

AND occurrence.hasgeospatialissues = FALSE

AND NOT GBIF_STRINGARRAYCONTAINS(occurrence.issue, 'TAXON_MATCH_FUZZY', TRUE)

AND (occurrence.distancefromcentroidinmeters >= 2000.0 OR occurrence.distancefromcentroidinmeters IS NULL)

AND (occurrence.specieskey IS NOT NULL AND occurrence."year" IS NOT NULL AND occurrence."month" IS NOT NULL AND occurrence.hascoordinate = TRUE)

GROUP BY

occurrence.kingdom,

occurrence.kingdomkey,

occurrence.phylum,

occurrence.phylumkey,

occurrence.class,

occurrence.classkey,

occurrence."order",

occurrence.orderkey,

occurrence.family,

occurrence.familykey,

occurrence.genus,

occurrence.genuskey,

occurrence.species,

occurrence.specieskey,

PRINTF('%04d-%02d', occurrence."year", occurrence."month"),

GBIF_EQDGCODE(2, occurrence.decimallatitude, occurrence.decimallongitude, 0.0)



### References

[1] Desmet P (2024). Download a species occurrence cube from GBIF.org. <https://docs.b-cubed.eu/tutorials/download-a-cube-from-gbif/>

[2] GBIF (2025). <https://techdocs.gbif.org/en/data-use/api-sql-downloads>. Accessed on 26 August 2025
