## Appendix B for "b3alien: A Python package to assess the introduction rate of alien species in a FAIR and reproducible way"

APPENDIX B: Exemplar workflow to determine the change in Rate of Establishment

This Jupyter notebook will guide the user of the b3alien Python package to generate a measure of the rate of establishment of alien species in a specific region or country. This workflow can be modified to match the needs of the user.

An executable version of this Jupyter notebook can be found here: <https://github.com/mtrekels/b3alien/tests/notebooks/exemplar_workflow.ipynb>.

Instructions indicated with a red border are optional or you only need to run these steps in very specific cases (e.g. when a GeoParquet file doesn’t exist or the taxonomic matching of the GRIIS checklist needs to be performed)

### Install the b3alien Python package

pip install b3alien

### Load the software package and additional libraries

from b3alien import b3cube

from b3alien import simulation

from b3alien import griis

import pandas as pd

import geopandas as gpd

import folium

from folium import Choropleth

from IPython.display import display

import matplotlib.pyplot as plt

%matplotlib inline

### Step 1: Generate a Biodiversity Data Cube from GBIF

This step can be skipped if you already have generated a GeoParquet file with your data cube.

The basic workflow assumes as an input a basic occurrence cube generated on the GBIF infrastructure. It uses the SQL API interface to GBIF data. How to generate this cube is described in Appendix A.

As a result of the cube generation through GBIF, you will be able to download a CSV file, which only contains the cell code of the chosen geographic grid. In order to match the grid with a vector geometry, the b3alien package supports the generation of GeoParquet files, if you have a GeoPackage of your grid. In the rest of this notebook, we used the Extended Quarter Degree grid cells, for which you can find GeoPackage files here: <https://download.gbif.org/grids/EQDGC/> or on Zenodo.

Once you have both files, you can generate the GeoParquet file by:

from b3alien import utils

utils.to_geoparquet(csvFile, geoFile, leftID='eqdcellcode', rightID='cellCode', exportPath='./data/export.parquet')



In the above code block you need to replace the names of you locally saved files and the corresponding names of the columns in these files.

More information on this function can be found on the documentation website. However, if you don't need the geospatial mapping of your results, it is not needed to generate a GeoParquet file. The b3alien package also detects when a pure occurrence cube in CSV format is used.

### Step 2: Download the GRIIS checklist of the considered country, and load it as a checklist object

In this exemplar workflow, we will use the checklist of Costa Rica, which is made available in the test data directory of the GitHub repository. However, you can download any of the checklists from GBIF.

To ensure that a correct matching of the species with the GBIF taxonomic backbone is available, you should run the following code snippet in the checklist data (replace the path to the path of your checklist):

griis.do_taxon_matching('../data/costa-rica/dwca-griis-costa-rica-v1.4')



This function is calling the GBIF API. Be gentle with it, and don't use it too much in a short time frame.

Now an additional file is generated (merged_distr.txt), which can be loaded as a CheckList object

CL = griis.CheckList("../data/costa-rica/dwca-griis-costa-rica-v1.4/merged_distr.txt")

### Step 3: Load the OccurrenceCube

A locally pre-generated data cube can be loaded by calling the following function:

cube = b3cube.OccurrenceCube("../data/costa-rica/data_CR_level2.parquet")



In case you have a GeoParquet file stored on Google Cloud, you can also load the cube using the same function, but by creating the OccurrenceClass by providing the link to the file stored in the cloud, and the corresponding Google project. For example:

cube = b3cube.OccurrenceCube("gs://b-cubed-eu/data_BE.parquet", gproject='$YOUR_GPROJECT')



Support for other cloud providers might be implemented in a next version of the software.

Additionally, it is possible to add "Observed Species Richness" as a property of the data cube. However, this is not needed for the remainder of the workflow.

cube._species_richness()



With the built in plotting functions, it is possible to plot the observed speciies richness on a Folium map.

b3cube.plot_richness(cube)



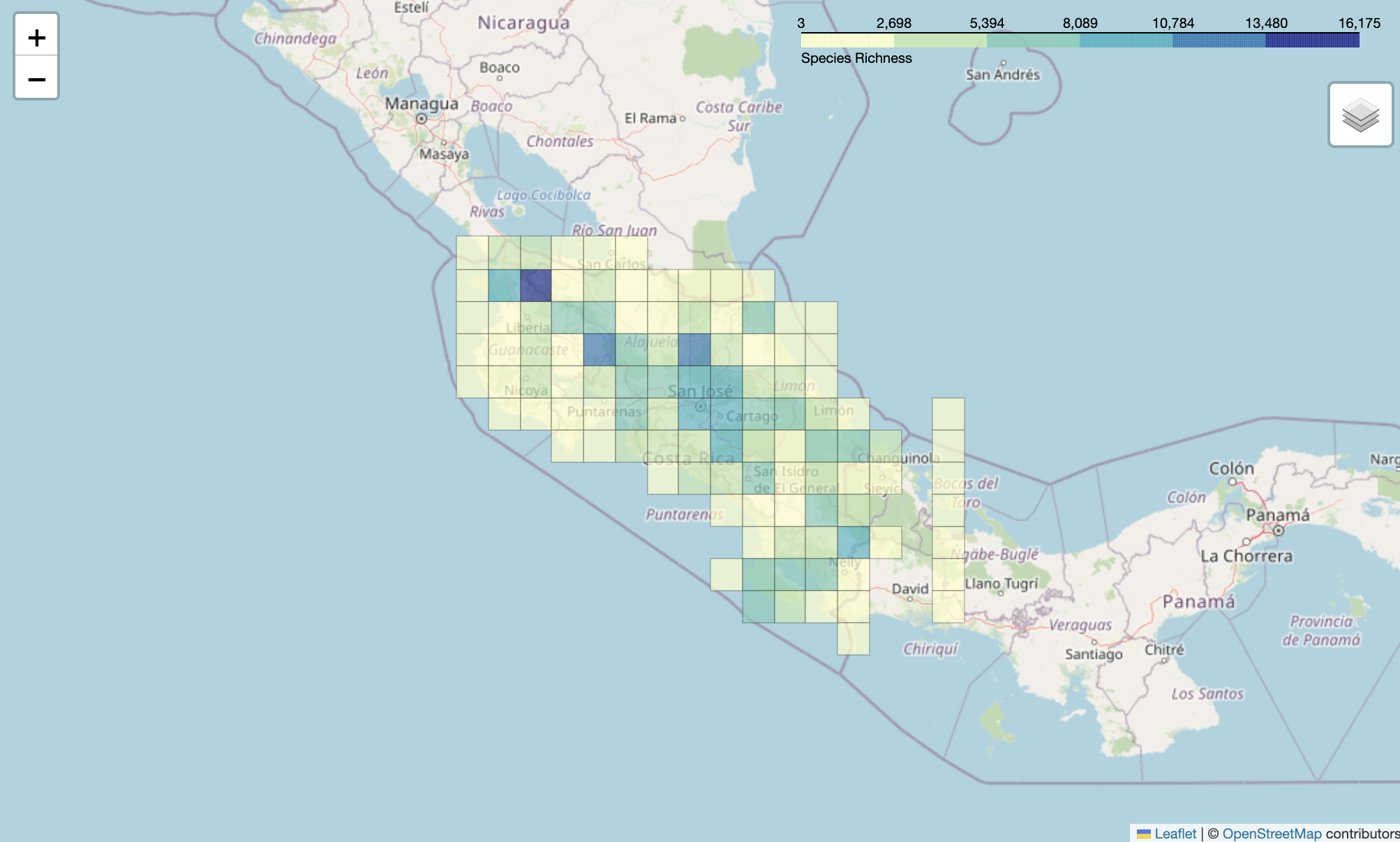

### Step 3: Calculate the Rate of Establishment from the CheckList and OccurrenceCube

After generating the basic objects, we can calculate the cumulative number of introduced species, and determine the rate of introduction by the Solow-Costello algorithm.

d_s, d_c = b3cube.cumulative_species(cube, CL.species)



This function is returning two dataframes, a sparse dataframe (d_s) which contains the cumulative number of species per cell, and a dataframe (d_c) which is cell independent. It is this last dataframe that we will use in the rest of the calculation. The first dataframe might be useful to have some spatial insight in the number of species.

From there, we can calculate the rate:

time, rate = b3cube.calculate_rate(d_c)



For applying the strategy defined by the GBF, is it useful to calculate the rate of introduction for different time windows. Therefore, a filtering function was developed to determine for which time interval the calculation needs to be done. In the rest of this exemplar notebook, we use the time frame 1970 - 2022.

df = pd.DataFrame({

```
    "year": time,
    "rate": rate
})

time, rate = b3cube.filter_time_window(df, 1970, 2022)
```



With the filtered time and rate, we can simulate the rate of introduction:



_, vec1 = simulation.simulate_solow_costello_scipy(time, rate, vis=True)



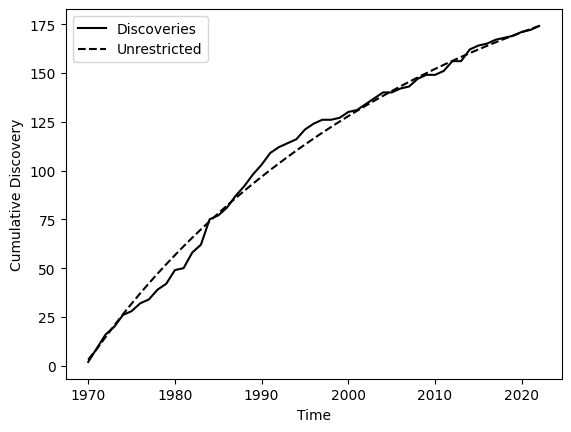

The vector 'vec1' contains the parameters of the fitting of the Solow-Costello model. The most important parameter in this case is the change in rate of establishment (2nd parameter).

print("Fitted Rate of Establishment from the data cube: " + str(vec1[1]) + "/year")



Fitted Rate of Establishment from the data cube: -0.025016351861057464/year

### Step 4: Determine the error margins on the fitted rate of establishment

The software package also includes the determination of the 95% confidence interval. This step is most efficiently run on a multi core machine, since it is using parallel computing. The method is using bootstrapping on the residuals.

results = simulation.parallel_bootstrap_solow_costello(time, rate, n_iterations=200)

simulation.plot_with_confidence(time, rate, results)



Bootstrapping: 100%|██████████| 200/200 [00:28<00:00, 6.90it/s]



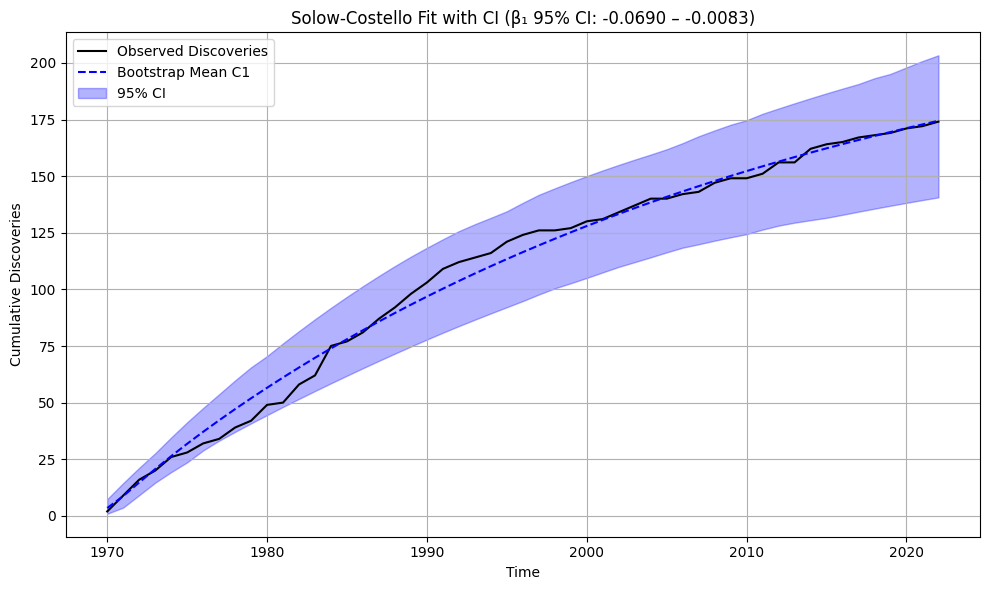

### Step 5: Determine the survey effort

In the reporting procedure for the GBF Target 6 indicator, there is a requirement of plotting the survey effort next to the cumulative discovery of species to have an idea on the detectability. In this section we give 2 examples of how you can approach this using the pure GBIF occurrence cube.

#### Aggregate the counts at a higher taxonomic level

One possible strategy is to count the total number of occurrences at a higher taxonomic level. In this example, we count the total number of counts for the kingdom Plantae per cell:

gdf = b3cube.aggregate_count_per_cell(cube, "kingdom", "Plantae")



Aggregation in space:

aggregated_gdf_space = gdf.groupby("cellCode").agg({

```
    "kingdomcount": "sum",
    "geometry": "first"  # assumes geometry is unique per cellCode
}).reset_index()
```



/var/folders/x8/_09h8jls5d3fclg4zvtq291c0000gn/T/ipykernel_54380/1479949437.py:1: FutureWarning: The default of observed=False is deprecated and will be changed to True in a future version of pandas. Pass observed=False to retain current behavior or observed=True to adopt the future default and silence this warning.

aggregated_gdf_space = gdf.groupby("cellCode").agg({



Aggregation in time:

aggregated_gdf_time = gdf.groupby("yearmonth").agg({

```
    "kingdomcount": "sum" # assumes geometry is unique per cellCode
}).reset_index()
```



/var/folders/x8/_09h8jls5d3fclg4zvtq291c0000gn/T/ipykernel_54380/1168206669.py:1: FutureWarning: The default of observed=False is deprecated and will be changed to True in a future version of pandas. Pass observed=False to retain current behavior or observed=True to adopt the future default and silence this warning.

aggregated_gdf_time = gdf.groupby("yearmonth").agg({



Afterwards you can plot for example the aggregated counts for one year, to get an idea on the survey effort per year:

aggregated_gdf_time = aggregated_gdf_time[aggregated_gdf_time['yearmonth'].str[:4].astype(int) >= 1677]

```
aggregated_gdf_time['yearmonth'] = pd.to_datetime(aggregated_gdf_time['yearmonth'], format='%Y-%m')
aggregated_gdf_time['year'] = aggregated_gdf_time['yearmonth'].dt.year

yearly_agg = aggregated_gdf_time.groupby('year')['kingdomcount'].sum().reset_index()
yearly_agg = yearly_agg.sort_values('year')


def filter_time_window(df, start_year, end_year):
    """Filter time and rate based on year window."""
    filtered = df[(df["year"] >= start_year) & (df["year"] <= end_year)].reset_index(drop=True)
    return filtered["year"], filtered["kingdomcount"]

time, obs = filter_time_window(yearly_agg, 1970, 2022)
```



plt.figure(figsize=(10, 5))

plt.plot(yearly_agg["year"], yearly_agg["kingdomcount"], marker="o")

plt.title("Obervation effort")

plt.xlabel("Time")

plt.ylabel("Observed Count")

plt.xticks(rotation=45)

plt.grid(True)

plt.tight_layout()

plt.show()



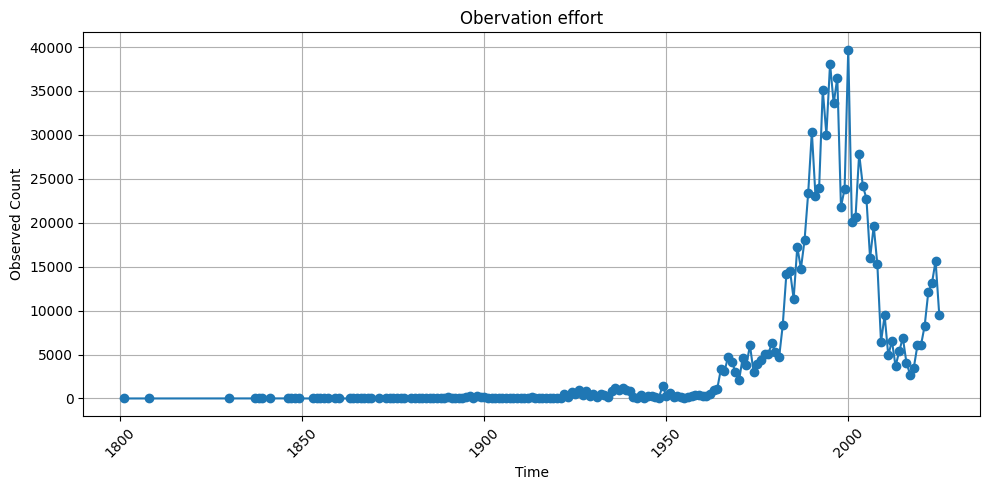

#### Survey effort by determining the number of distinct observers

This is another potential measure of the survey effort. If you created the data cube accorrding to Appendix A, this measure is readily available.

survey_eff = b3cube.get_survey_effort(cube, calc_type='distinct')



def filter_time_window(df, start_year, end_year):

```
    """Filter time and rate based on year window."""
    df["date"] = df["date"].dt.year.astype(int)
    filtered = df[(df["date"] >= start_year) & (df["date"] <= end_year)].reset_index(drop=True)
    return filtered["date"], filtered["distinct_observers"]

time2, observ = filter_time_window(survey_eff, 1970, 2022)
```



Using the results from the simulation in previous steps, we can now plot the survey effort and the cumulative distribution together on a single plot. For clarity, we plot the logaritmn of the survey effort, since there is a huge amount of GBIF data that is coming from citizen science programs. This results in an enormous increase in available data.

import numpy as np

```
# Create a figure and the first set of axes (ax1)
# This is the object-oriented way, which is better for complex plots.
fig, ax1 = plt.subplots(figsize=(12, 6))

# --- Plotting the first dataset (left y-axis) ---
color = 'tab:blue'
ax1.set_xlabel('Time')
ax1.set_ylabel('Log(Observation effort)', color=color) # Label for the left y-axis
ax1.plot(time2, np.log(observ), marker='o', linestyle='-', color=color, label='Observation effort')
ax1.tick_params(axis='y', labelcolor=color)
ax1.grid(True) # Add grid lines

# --- Creating and plotting on the second y-axis ---
# Create a second axes that shares the same x-axis
ax2 = ax1.twinx()

color = 'tab:red'
ax2.plot(time, np.cumsum(rate), 'k-', label='Observed Discoveries')
ax2.plot(time, results["c1_mean"], 'b--', label='Bootstrap Mean C1')
ax2.fill_between(time, results["c1_lower"], results["c1_upper"], color='blue', alpha=0.3, label='95% CI')
ax2.set_ylabel('Cumulative Discoveries')
ax2.tick_params(axis='y', labelcolor=color)

# --- Final Touches ---
plt.title('Solow-Costello fit and Observation Effort')
fig.autofmt_xdate(rotation=45) # A better way to handle date rotation

# To create a single legend for both lines
# Get handles and labels from both axes and combine them
lines1, labels1 = ax1.get_legend_handles_labels()
lines2, labels2 = ax2.get_legend_handles_labels()
ax2.legend(lines1 + lines2, labels1 + labels2, loc='upper left')

# Ensure everything fits without overlapping
fig.tight_layout()

plt.show()
```



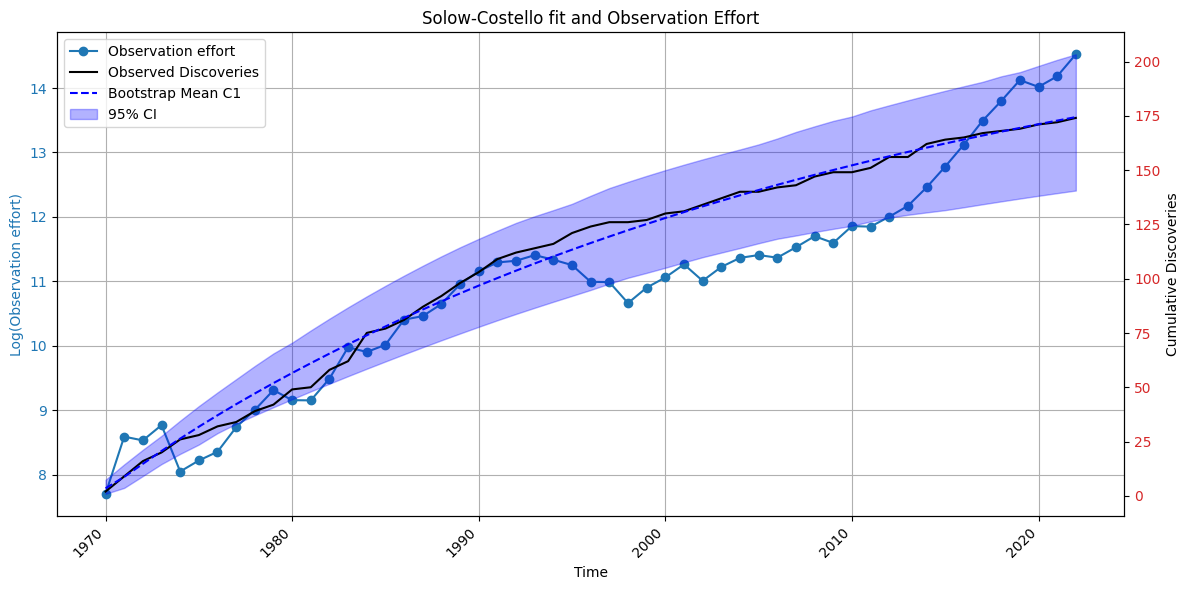

To get an idea on the geographical distribution of the survey effort, we can also plot this on a map. For example for the aggregates counts at kingdom level, we can have the following results:

survey_eff_spatial = gpd.GeoDataFrame(aggregated_gdf_space, geometry="geometry", crs=gdf.crs)



survey_eff_spatial["geometry"] = survey_eff_spatial["geometry"].simplify(0.01)

```
centroid = survey_eff_spatial.geometry.unary_union.centroid
m = folium.Map(location=[centroid.y, centroid.x], zoom_start=7)

# Plot
Choropleth(
    geo_data=survey_eff_spatial,
    name='Aggregated',
    data=survey_eff_spatial,
    columns=['cellCode', 'kingdomcount'],
    key_on='feature.properties.cellCode',
    fill_color='YlGnBu',
    fill_opacity=0.7,
    line_opacity=0.2,
    legend_name='Observation effort'
).add_to(m)

folium.LayerControl().add_to(m)
```



/var/folders/x8/_09h8jls5d3fclg4zvtq291c0000gn/T/ipykernel_54380/20851828.py:2: DeprecationWarning: The 'unary_union' attribute is deprecated, use the 'union_all()' method instead.

```
  centroid = survey_eff_spatial.geometry.unary_union.centroid


<folium.map.LayerControl at 0x3d0280e30>
```



display(m)



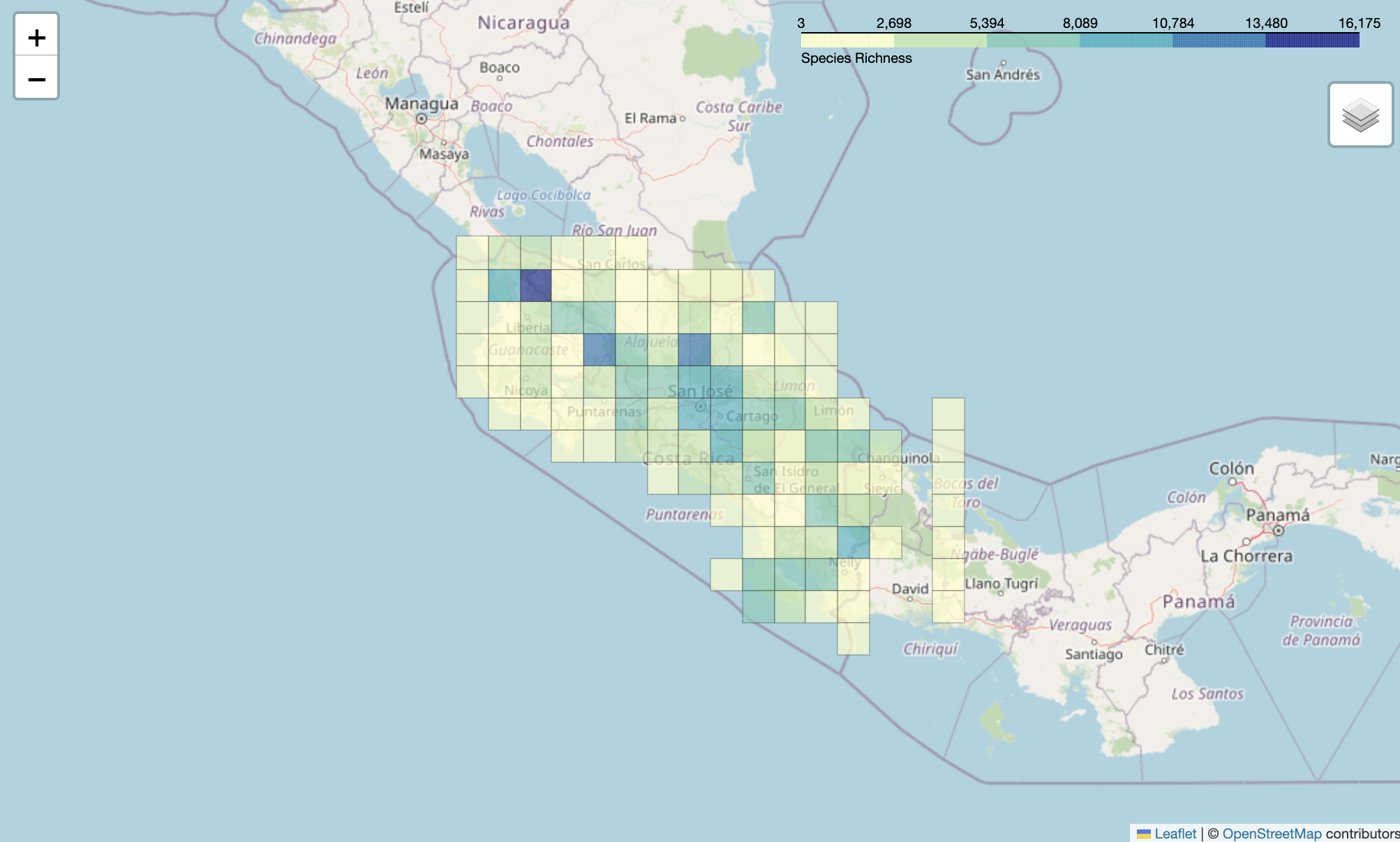

#### Additional example: plotting of the number of distinct observers per grid cell

aggregated_obs_space = cube.df.groupby("cellCode").agg({

```
    "distinctobservers": "sum",
    "geometry": "first"  # assumes geometry is unique per cellCode
}).reset_index()
```



/var/folders/x8/_09h8jls5d3fclg4zvtq291c0000gn/T/ipykernel_54380/1800479429.py:1: FutureWarning: The default of observed=False is deprecated and will be changed to True in a future version of pandas. Pass observed=False to retain current behavior or observed=True to adopt the future default and silence this warning.

aggregated_obs_space = cube.df.groupby("cellCode").agg({



obs_space = gpd.GeoDataFrame(aggregated_obs_space, geometry="geometry", crs=cube.df.crs)



gdf.crs

```
obs_space["geometry"] = obs_space["geometry"].simplify(0.01)
centroid = gdf.geometry.unary_union.centroid
m = folium.Map(location=[centroid.y, centroid.x], zoom_start=7)

# Plot
Choropleth(
    geo_data=obs_space,
    name='Aggregated',
    data=obs_space,
    columns=['cellCode', 'distinctobservers'],
    key_on='feature.properties.cellCode',
    fill_color='YlGnBu',
    fill_opacity=0.7,
    line_opacity=0.2,
    legend_name='Observation effort'
).add_to(m)

folium.LayerControl().add_to(m)

display(m)
```



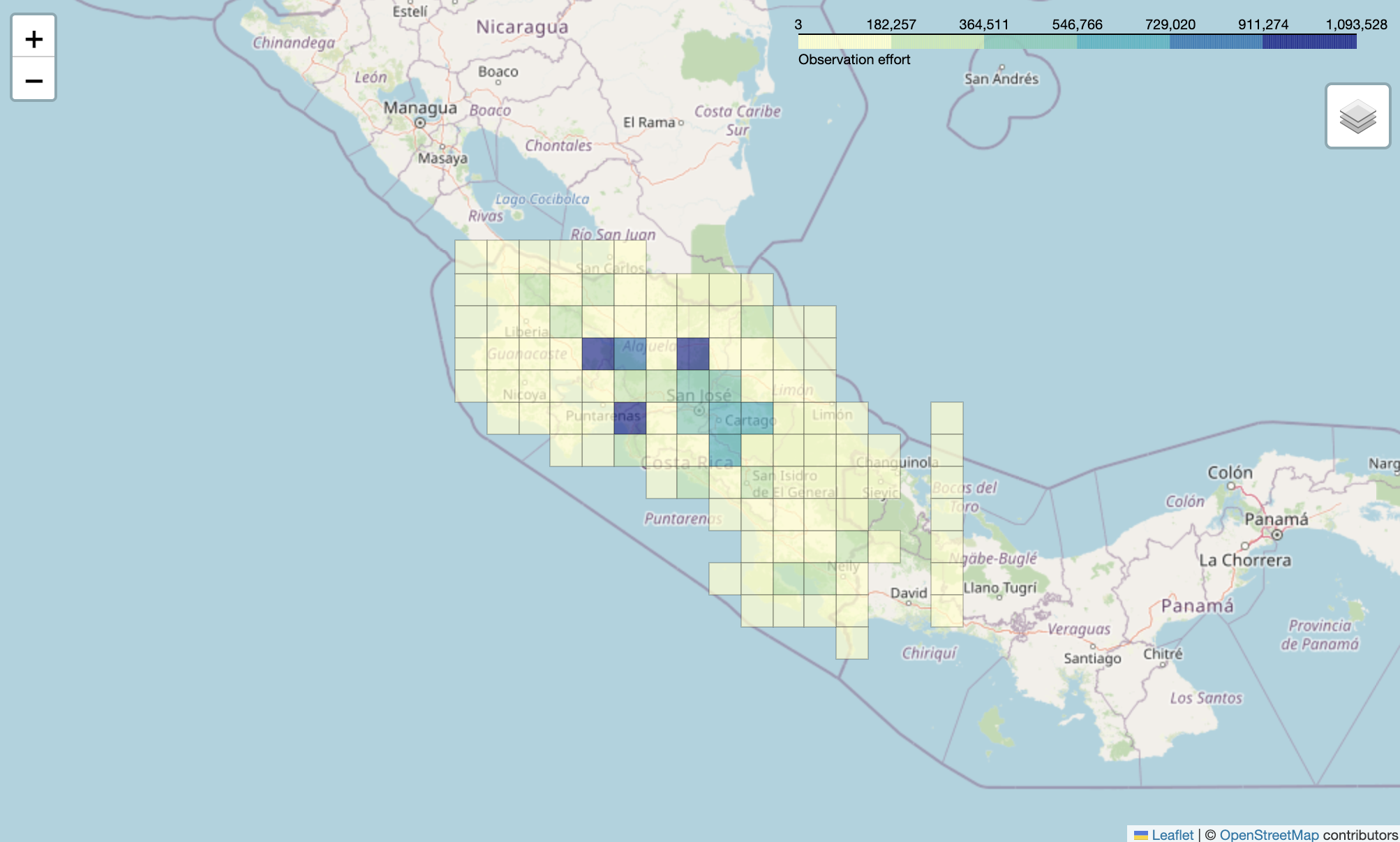
